# Evidence of a recombination rate valley in human regulatory domains

**DOI:** 10.1101/048827

**Authors:** Yaping Liu, Abhishek Sarkar, Manolis Kellis

## Abstract

Human recombination rate varies greatly, but the forces shaping it remain incompletely understood. Here, we study the relationship between recombination rate and gene-regulatory domains defined by a gene and its linked control elements. We define these links using methylation quantitative trait loci (meQTLs), expression quantitative trait loci (eQTLs), chromatin conformation, and correlated activity across cell types. Each link type shows a “recombination valley” of significantly-reduced recombination rate compared to control regions, indicating preferential co-inheritance of genes and linked regulatory elements as a single unit. This recombination valley is most pronounced for gene-regulatory domains of early embryonic developmental genes, housekeeping genes, and constitutive regulatory elements, which are known to show increased evolutionary constraint across species. Recombination valleys show increased DNA methylation, reduced double-stranded break initiation, and increased repair efficiency, specifically in the lineage leading to the germ line, providing a potential molecular mechanism facilitating their maintenance by exclusion of recombination events.

## Introduction

Variation in recombination rates in humans and other diploid organisms can be shaped by both evolutionary and molecular processes(Coop and Przeworski 2007), but these are only partially understood. High-resolution human recombination maps have been estimated using both parent-offspring transmission(Kong et al. 2002; Kong et al. 2010) and patterns of linkage disequilibrium (LD)(McVean et al. 2004; Hinds et al. 2005; Myers et al. 2005; Consortium et al. 2010). These have revealed highly localized regions with higher or lower recombination rate, known as recombination hotspots and coldspots, respectively(Myers et al. 2005). Sequences analysis showed that human recombination hotspots are associated with a number of sequence features such as PRDM9 binding motifs(Baudat et al. 2010), CpG islands, and GC-rich repeats(Myers et al. 2005; Consortium et al. 2010; Auton et al. 2012), and that recombination coldspots are associated with repetitive elements, transcribed regions, and telomeres(McVean et al. 2004; Myers et al. 2005).

Outside recombination hotspots, differences in epigenomic signatures are associated with differences in recombination rate(Sigurdsson et al. 2009; Zeng and Yi 2014). In particular, the level of DNA methylation, which is primarily established at prophase I when recombination occurs(Oakes et al. 2007), is reported to be positively correlated with recombination rate (Zeng and Yi 2014). A causal effect of DNA methylation on recombination rate was established using a methylation-deficient strain of *Arabidopsis*, which showed reduction of recombination rate in euchromatic regions(Melamed-Bessudo and Levy 2012; Mirouze et al. 2012).

## Results

### Gene-regulatory domains defined using eQTLs and meQTLs show a recombination valley

We first studied the relationship between human recombination rate and regulatory domains, defined as the genomic region spanned by a gene and all the regulatory regions linked to its promoter element. We estimated recombination rates using the 1000 Genomes genetic map(Consortium et al. 2010), and related them to gene-regulatory domains using four types of links.

We first used genetic links based on expression Quantitative Trait Loci (eQTLs) and methylation Quantitative Trait Loci (meQTLs). These consist of 248,856 eQTL links between regulatory regions and their target genes’ Transcription Starting Sites (TSS), defined in whole blood using 168 individuals profiled by the Gene-Tissue Expression (GTEx) Consortium(Consortium 2013), and 809,577 meQTL links between regulatory regions and CpG methylation measured in human brain primarily in promoter regions in the ROS/MAP cohort of 575 individuals(De Jager et al. 2014).

We found that the intervals between eQTLs and their target genes and between meQTLs and their target methylation probes are substantially depleted for recombination hotspots (Fig. 1a-b). We evaluated intervals at three distance ranges, consisting of short (1kb-10kb), intermediate (10kb-100kb) and long (100kb-1Mb) distances. The effect was most pronounced for links of intermediate and long distances, which showed consistently lower recombination rates compared to random intervals, a phenomenon which we call a “recombination valley” (Fig. 1e-f; Supplementary Fig. 1a-b). To evaluate the statistical significance of this observation, we sampled the same number of random genomic regions with the same physical length in the same chromosome (Supplemental method 1). We also sampled the same number of random SNP-TSS/SNP-CpG pairs as another set of random intervals (Supplemental method 2). Comparing with these two random datasets, we found a significant decrease in both cases (One-way paired Mann-Whitney U test and permutation test, p<1e^−4^) in recombination rate at both intermediate and long distances and for both eQTL and meQTL links (Fig. 1i-j).

**Figure 1.**
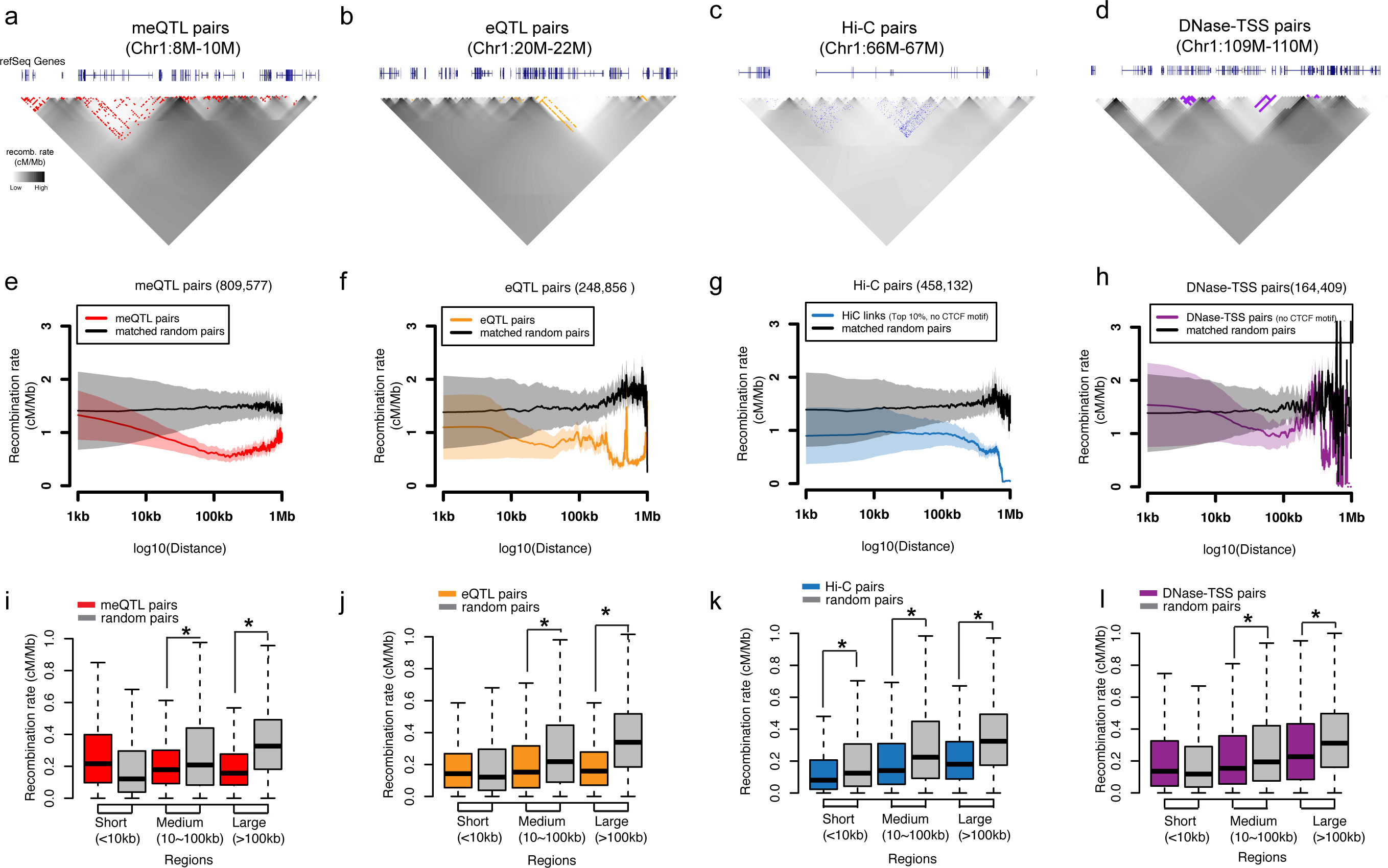
Recombination valley within genetics, physical and activity links. Individual genomic regions about enrichment of (a) meQTL pairs (red), (b) eQTL pairs(orange), (c) top 10 % Hi-C pairs (Observed/Expectation, no CTCF motif, blue), (d) DNase-TSS pairs (no CTCF motif, purple) in low recombination rate regions. Each pixel represents 10kb segment (for Hi-C, it represent 1kb segment), the more dark color indicates higher recombination rate between two segments. Colored dot at each pixel indicates the genetics or physical links exist between two genomic segments. The average recombination rate within (e) meQTL pairs (red), (f) eQTL pairs (orange), (g) top 10% Hi-C pairs (Observed/Expectation, no CTCF motif, blue), (h) DNase-TSS pairs (no CTCF motif, purple) are significantly lower than the matched random intervals. X-axis is log10 base pair distance between linked genomic features. Y-axis is the average recombination rate within each interval. Colored lines represent the mean recombination rate at each interval distance between genomic features, while black links represent the mean value in matched random intervals. Shaded regions represent the 95% confident interval. Boxplot of the recombination rate in three different genomic scales in (i) meQTL pairs, (j) eQTL pairs, (k) top 10% Hi-C pairs (Observed/Expectation, no CTCF motif), (l) DNase-TSS pairs (no CTCF motif). One-way paired Mann-Whitney U test show that recombination rate within genetics, physical and activity links at medium and large intervals is significantly less than random matched pairs. Comparisons with p<1e-4 are marked with *.

We next confirmed that our observation in genetics-based links is not an artifact from Linkage Disequilibrium (LD). First, we only keep the best eQTL/meQTL with the highest false discovery rate (FDR) for each gene/CpG. We further pruned the best eQTL /meQTL links by excluding the multiple counts of the same genomic region (Supplemental method 3) and consistently found a significant recombination valley in the regions defined by genetic links (Supplementary Fig. 2a-c, f-h, k-m). Second, we made a small random shift around the best genetic links and found slightly but significantly higher recombination rate (Supplemental method 3, Supplementary Fig. 3). This observation suggests that causal genetics links showed lowest recombination rate within LD.

We further evaluated whether our observation also held in independent datasets and with varying analytical parameters. First, we repeated this analysis using 16 additional tissues and cell lines from the GTEx Consortium(Consortium 2013), the Multiple Tissue Human Expression Resource (MuTHER) Consortium(Grundberg et al. 2012) and Genetic European Variation in Health and Disease (gEUVADIS) Consortium(Lappalainen et al. 2013) and consistently found a significant recombination valley in the regions defined by genetic links (Supplementary Fig. 4). Second, we repeated the analysis varying thresholds for the false discovery rate (FDR) of eQTLs and meQTLs and consistently found a recombination valley. The strongest reduction in recombination rate was found at the most stringent FDR thresholds for eQTL and meQTL discovery, indicating that with higher link confidence thresholds, the signal becomes stronger (Supplementary Fig. 5a-b, e-f). Third, we repeated the analysis on genetic maps estimated by the HapMap project(International HapMap et al. 2010) and by deCODE Genetics(Kong et al. 2002; Kong et al. 2010) and found the results are largely unchanged (Supplementary Fig. 6a-b,e-f). To account for sequence biases, we additionally generated matched random intervals, which have equal GC content, CpG density, SNP density, and PRDM9 motif density. We found the results are robust to more stringent matching (Supplementary Fig. 7ab, e-f, i-j, m-n, q-r). To account for the decreased recombination rate in transcribed regions, we excluded the intervals within 2kb of gene annotations in GENCODE v19 and still found recombination valleys in meQTL/eQTL links comparing with random intervals that were generated with more stringent matched criteria (Supplementary Fig. 8a-b, e-f).

### Gene-regulatory domains by chromosome conformation show a recombination valley

In addition to genetic links, we used 458,132 links between genomic regions that are in close proximity when folded in the three-dimensional nucleus, based on chromosomal interactions from high-throughput chromosome conformation capture (Hi-C) measured in the GM12878 cell line(Rao et al. 2014). We found that the recombination rate within regulatory domains defined by Hi-C was also significantly lower (Oneway paired Mann-Whitney U test and permutation test, p<1e^−4^) at both intermediate and long distances comparing with two different sets of random intervals (Fig. 1c,g,k, Supplementary Fig. 1c). This property held specifically for Hi-C links that are not interrupted by CTCF motifs, consistent with the role of CTCF loops as defining regulatory domain boundaries(Rao et al. 2014) (Fig. 1g,k, Supplementary Fig. 11o,p).

To avoid the bias introduced by relatively more Hi-C links from the domains with lower recombination rate, we generated the matched random intervals only within the same HiCCUPS loops and still found recombination valley (Supplementary Fig. 9). We also pruned the Hi-C links by excluding the multiple counts of the same genomic region and consistently found recombination valley (Supplementary method 3. Supplementary Fig. 2d, i, n). We next varied the threshold for Hi-C links included in the analysis and observed recombination valley again in different threshold choices (Supplementary Fig. 5c,g). We also repeated the analysis in different genetic maps (Supplementary Fig 6c,g) and compared it with more stringent matched random intervals in whole genome (Supplementary Fig. c,g,k,o,s) and non-coding regions (Supplementary Fig 8c,g). We also combined Hi-C and eQTL evidence available in the same cell type (lymphoblastoid cell lines, LCLs, including GM12878)(Grundberg et al. 2012; Rao et al. 2014), and found that the depletion in the recombination rate became even more pronounced (Supplementary Fig. 10). This indicates that gene-regulatory links with increased confidence show an even more pronounced recombination valley.

We repeated this analysis using physical chromosomal interactions defined using Chromatin Interaction Analysis using Paired-End-Tag sequencing (ChIA-PET), a complementary technique that defines long-range looping interactions for regions that interact in the context of a specific regulator. We used regulatory domains based on ChIA-PET for both polymerase PolII and CTCF, as defined by the ENCODE consortium. We found that the recombination rate within ChIA-PET PolII domains was also significantly depleted (Supplementary Fig. 11k-n).

These results indicate that the recombination valley is a general property of gene-regulatory domains defined using long-range physical DNA interactions.

### Gene-regulatory domains defined using activity correlation show a recombination valley

We next evaluated the relationship between recombination rate and gene-regulatory links defined between enhancer regions and their target genes as predicted by the ENCODE(Ernst et al. 2011) and Roadmap Epigenomics Consortia(Roadmap Epigenomics et al. 2015). We used 29,557,079 unique correlation-based links predicted between DNase-seq peaks at enhancer chromatin states and DNase-seq peaks at the TSS of putative target transcripts (Kheradpour & Kellis. *In preparation*). Given the role of CTCF motifs in guiding chromatin loops(Rao et al. 2014), we focused on 164,409 unique links that were not interrupted by CTCF motifs and thus more likely to lie in the same chromatin loops. We found significantly reduced recombination rate for regions within these enhancer-gene domains relative to random pairs (Oneway paired Mann-Whitney U test and permutation test, p<1e^−4^, Fig. 1d, h, l, Supplementary Fig. 1d, Supplementary Fig. 11i-j).

To avoid the multiple counts from the same genomic region, we also pruned the DNase-TSS links by using the similar approach as in Hi-C links (Supplemental method 3) and found similar observations (Supplementary Fig. 2e, j, o). We next repeated the analysis by different thresholds for DNase-TSS links (Supplementary Fig 5d-h), different genetic maps (Supplementary Fig 6d-h) and more stringent matched random intervals in both of whole genome (Supplementary Fig 7d,h,l,p,t) and non-coding regions (Supplementary Fig 8d,h), and consistently found significantly reduced recombination valley within DNase-TSS links.

We did additional analysis with 1,427,744 unique enhancer-gene links predicted by a previously published algorithm(Ernst et al. 2011) based on the correlation between enhancer chromatin state calls and gene expression levels (Roadmap Epigenomics et al. 2015). We found that the resulting 139,043 gene-regulatory domains without CTCF motif continue to show a significant depletion in recombination rate at both intermediate and long distances (Supplementary Fig. 11a-d)

We also repeated this analysis using 302,538 unique links predicted using a module-based joint linear discriminant analysis (joint-LDA) linking approach (Wang & Kellis. *In preparation*) that does not depend on correlation, and can also predict cell type specific links. Despite these differences in predicting enhancer-gene links, we found a similar depletion in the recombination rate within gene-regulatory domains compared to random controls (Supplementary Fig. 11e-h).

Taken together, these results indicate that gene-regulatory domains defined based on functional genomics and epigenomic information are associated with a recombination valley, indicating that genes tend to be co-inherited with their gene-regulatory elements.

### Constitutive and developmental domains show stronger recombination rate depletion

We next evaluated how the strength of the recombination valley varies for different classes of genes. We found that the recombination valley within physical and activity links was more pronounced for housekeeping genes(Eisenberg and Levanon 2013) compared with non-housekeeping genes (Fig. 2a-f, Supplementary Fig. 12). It was also more pronounced for genes that act in early embryonic development stages(Xue et al. 2013; Guo et al. 2015), especially for genes in oocyte stage and genes responsible for meiosis in primordial germ cells (PGCs) stage, but not for most of other cell type-specific gene groups(Xie et al. 2013) (Supplementary Fig. 13).

**Figure 2.**
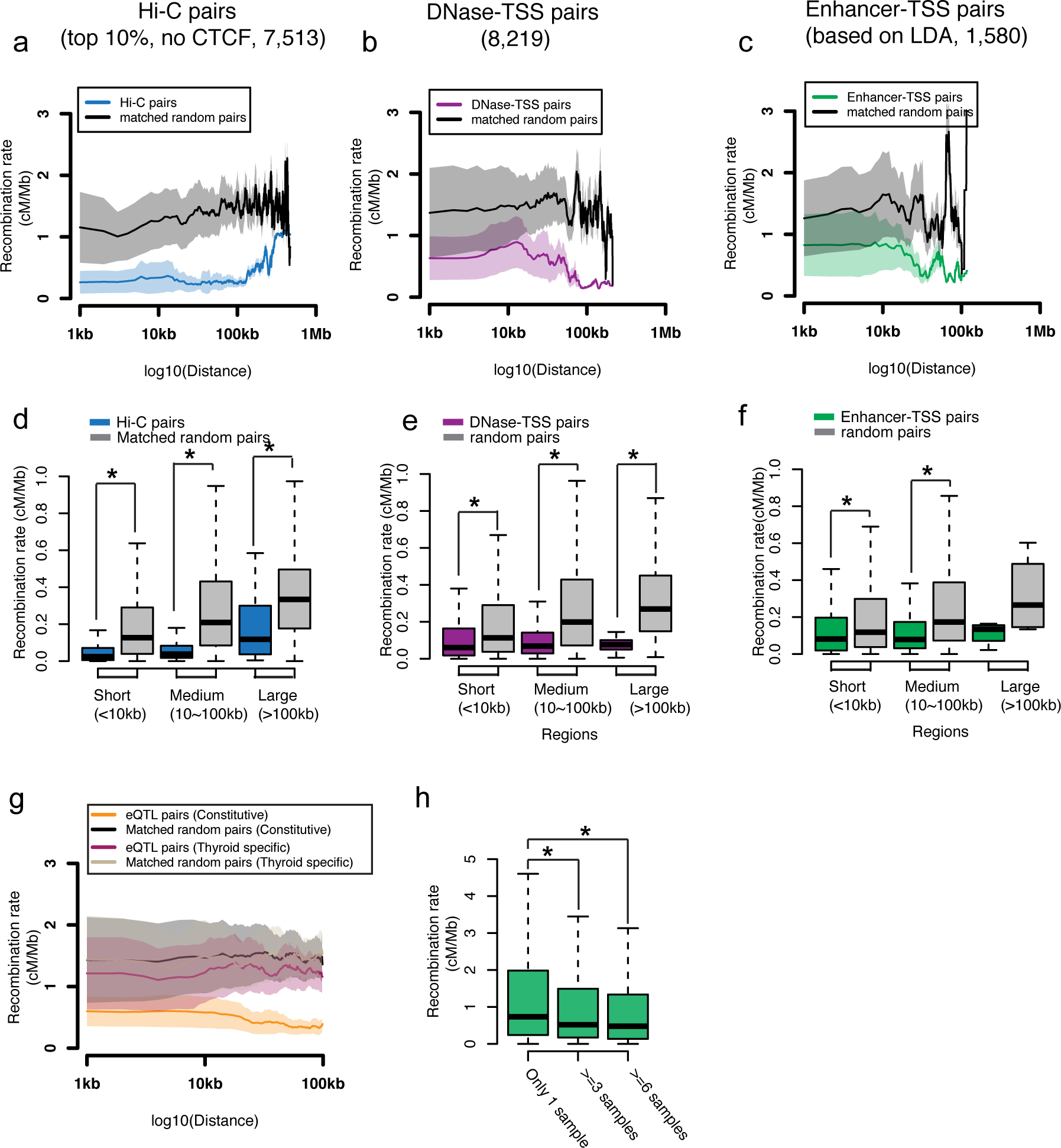
Recombination valley is most prominent at functional links associated with house keeping genes and constitutive links. (a) Average plot of recombination rate in top 10% Hi-C pairs (O/E, no CTCF) associated with house keeping genes (b) Average plot of recombination rate in DNase-TSS pairs without CTCF motif in-between that associated with house keeping genes. (c) Average plot of recombination rate in Enhancer-TSS pairs called by LDA method without CTCF motif in-between that associated with house keeping genes. (d) Box plot of recombination rate in top 10% Hi-C pairs (O/E, no CTCF) associated with house keeping genes. (e) Box plot of recombination rate in DNase-TSS pairs without CTCF motif in-between that associated with house keeping genes. (f) Box plot of recombination rate in Enhancer-TSS pairs called by LDA method without CTCF motif in-between that associated with house keeping genes. (g) Recombination valley is much more significant at constitute eQTL links (orange) than that at tissue specific eQTL links (magenta). Shaded regions represent the 95% confident interval. (h) Box plot of recombination rate within enhancer-TSS links (10kb-100kb region) called by joint-LDA method at different number of cell types.

We also evaluated how the strength of the recombination valley varies for different classes of regulatory elements. We used constitutive and tissue-specific eQTL links from the GTEx project, and evaluated the recombination rate depletion in regulatory domains defined by 43,236 constitutive eQTLs and 18,879 thyroid specific eQTLs(Consortium 2013). We found that constitutive eQTL links showed consistently larger discrepancies in recombination rates than tissue-specific links, each compared to matched random controls (Fig. 2g). Similarly, we found that gene-regulatory domains recovered independently in multiple cell types showed a more pronounced recombination valley than tissue-specific gene-regulatory domains (Fig. 2h).

Thus, the recombination valley is more strongly pronounced in gene-regulatory domains of constitutively-expressed genes, genes with developmental roles, and regulatory elements with constitutive activity, which all share the feature that they are under stronger evolutionary constraint. This suggests that a reduced recombination rate between regulatory elements and their target genes may be advantageous for genes and regulatory elements under stronger selection in the germ line lineage, possibly by facilitating maintenance of a gene and its regulatory elements as a single unit of inheritance during the early development.

### Recombination valley also found in mouse genetic data using mouse regulatory domains

We reasoned that if the recombination valley is a selected feature of gene-regulatory domains in human, it may be an evolutionarily conserved feature also found in other mammals. To test this hypothesis, we repeated our analysis in the mouse genome.

We quantified recombination rate across the mouse genome using the *Mus musculus* genetic map(Brunschwig et al. 2012). We defined gene-regulatory domains using both genetic and physical interactions. For genetic interactions, we used 2,659 eQTLs based on 100 strains in murine liver(Bennett et al. 2010) and 1,035 eQTLs based on 39 strains in two murine immunological cell types(Mostafavi et al. 2014). For physical interactions, we used 271,236 Hi-C links(Rao et al. 2014) called in a murine lymphoblastoid cell line.

Evaluating the recombination rate of gene-regulatory domains, we found a significant depletion in the recombination rate within domains relative to random pairs for both genetic and physical interactions (One-way paired Mann-Whitney U test and permutation test, p<1e^−4^, Fig. 3, Supplementary Fig. 14). This indicates that the recombination valley is not a feature solely of the human genome, but may represent a more general mammalian property, possibly as an evolutionary-conserved mechanism to preserve important regulatory domains.

**Figure 3.**
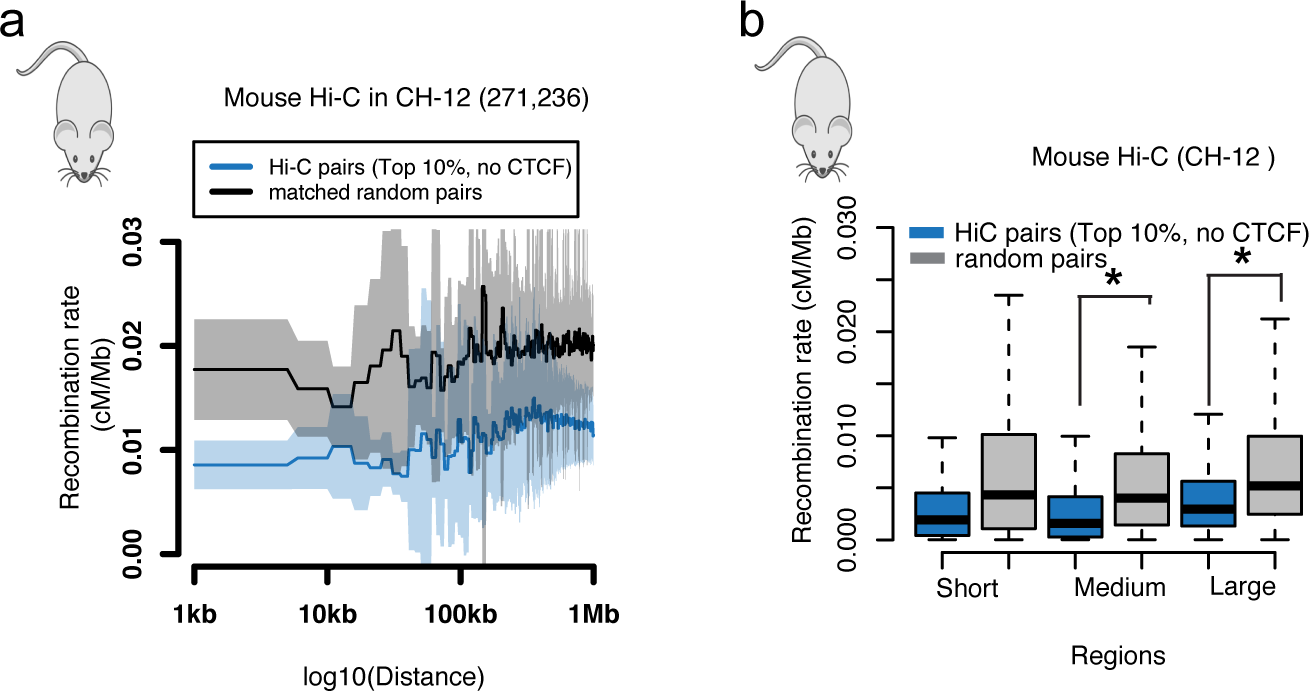
Recombination valley in mouse regulatory domains. (a) Average plot of recombination rate in top 10% Hi-C links (O/E) without CTCF peaks in-between at CH-12 cell. (b) Box plot of recombination rate in top 10% Hi-C links (O/E) without CTCF peaks in-between at CH-12 cell.

### Potential roles of DNA methylation and double-stranded breaks in recombination valley

We next sought to understand potential mechanistic processes that could lead to the observed recombination valley. As expected, we found that indeed a scarcity of recombination hotspots is associated with recombination valleys (Fig. 4a). This was accompanied by a reduction in the frequency of predicted PRDM9 binding events, indicating a potential role of PRDM9 exclusion in mediating the reduced recombination rate. However, half of links do not have recombination hotspots, and thus their recombination rate variation must be explained using other mechanisms.

**Figure 4.**
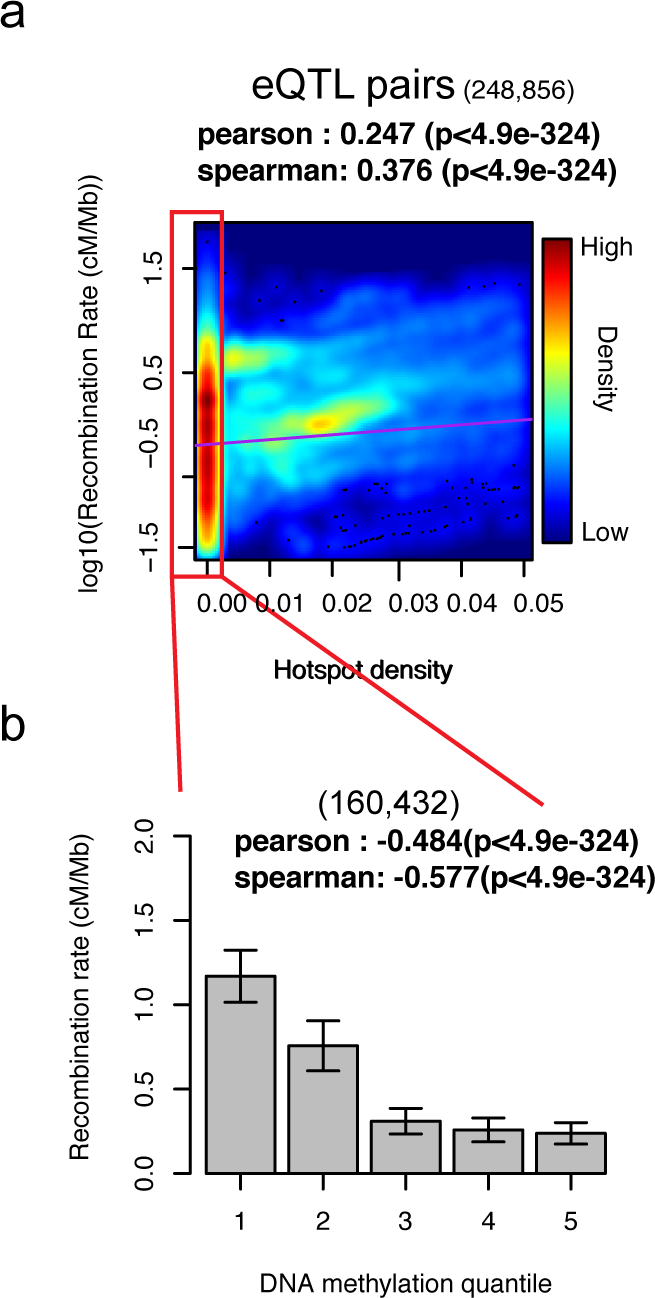
Recombination valley is correlated with hotspot density and DNA methylation. (a) Scatterplot for the relationship between log10(recombination rate) and recombination hotspot density (per kb) at each eQTL interval. Heat colors represent the point density. Recombination hotspot density less than 0.005/kb was circled by red rectangle and extracted out for the analysis in (b). (b) Barplot for the relationship between average recombination rate and DNA methylation quantiles in GV oocyte stage within eQTL intervals. Error bar indicates the standard deviation.

Given the previously proposed roles of DNA methylation in recombination rate, we studied the relationship between DNA methylation and recombination valleys. We used nucleotide-resolution genome-wide methylation profiles in human primordial germ cells (PGCs)(Guo et al. 2015) and oocytes(Okae et al. 2014), representing the methylome state of human cells both before and during meiotic arrest, in which recombination occurs via crossover events.

We used 500kb non-overlapping windows to scan the genome, and found a strong global negative correlation between methylation levels in PGCs and recombination rate (Fig. 5a-b), indicating that DNA methylation levels immediately prior to recombination events are highly predictive of recombination valleys. In contrast, we did not find this global anti-correlation in oocytes (Fig. 5b); however, methylation level within genetics links, such as eQTL links, in oocytes showed negative correlation with recombination rate (Fig. 4b, Figure 5c). These results suggest that DNA methylation may play a role in reducing the frequency of meiotic recombination events that disrupt functional regulatory links. We did not find strong global or local negative correlations between methylation and recombination rate in additional cell types in a number of developmental stages (Fig. 5c).

**Figure 5.**
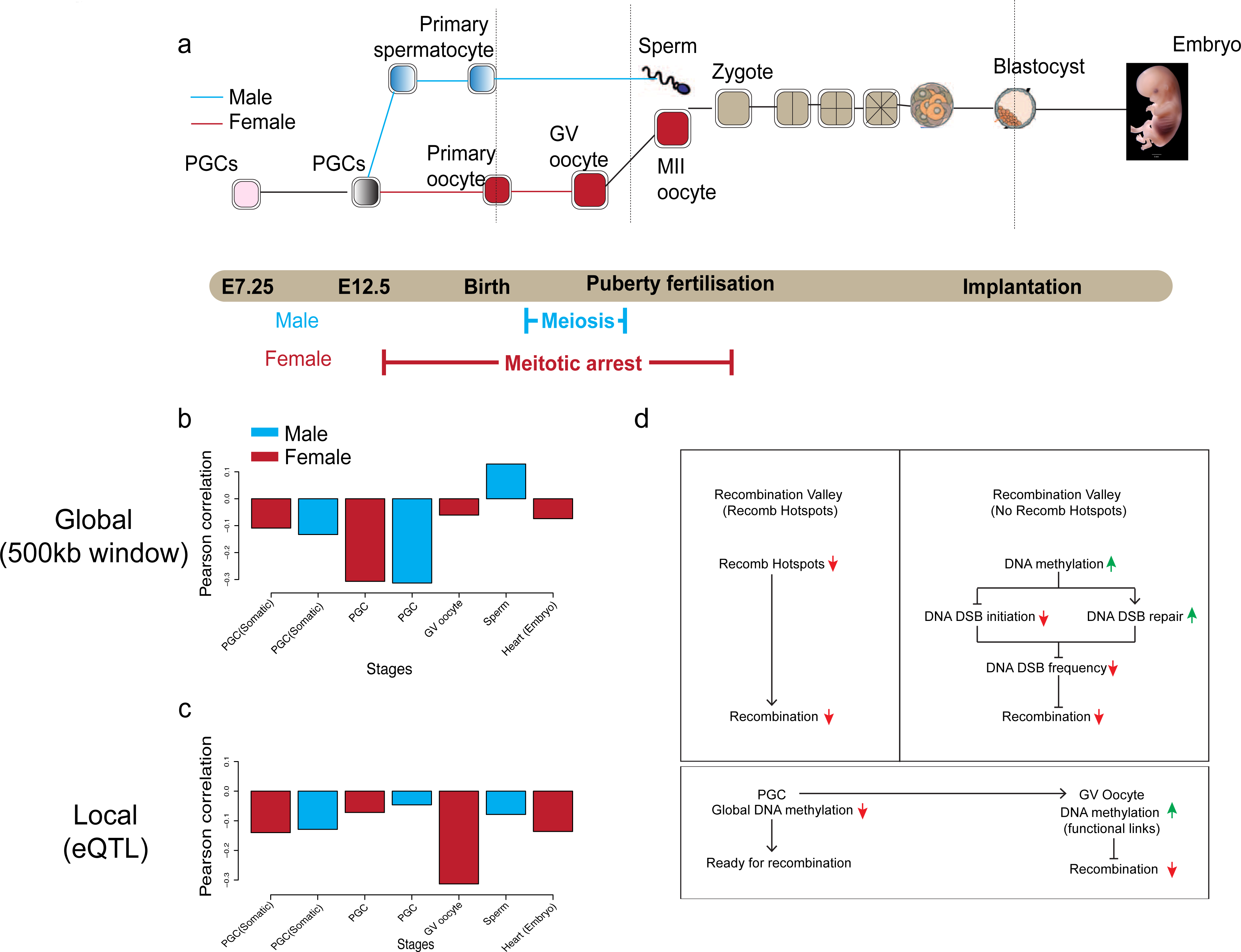
Relationship between recombination rate and DNA methylation quantile within 500kb window and within genetic links at different early development stages. (a) Flowchart of early development process in male and female. (b) Barplot of pearson correlation coefficient between recombination rate and DNA methylation quantiles within 500kb non-overlapped windows at different early development stages. (c) Barplot of pearson correlation coefficient between recombination rate and DNA methylation quantiles within eQTL pairs at different early development stages. Blue represents relationship in male, while red represents relationship in female. (d) The model of recombination valley formation at regions with and without recombination hotspot regions. In the region without recombination hotspot, the recombination valley formation at regulatory domain is associated with DNA methylation level at GV oocyte stage.

In absence of recombination hotspots (that are often mediated by PRDM9), double-stranded breaks can facilitate recombination. To quantify the combined predictive power of methylation levels and regulatory domains defined by genetic, physical and activity links, we built a random forest regression model that utilizes them as features to predict recombination rate, adjusting for the varying recombination rate of different chromosomes and at different genomic distances (see Supplementary method 12, 13).

We found that the individual link types varied greatly in their predictive power, with meQTLs showing the strongest predictive power for both medium-range and long-range links. We found that the combination of all four features resulted in high concordance between predicted and observed recombination rate (Pearson correlation coefficient 0.622, p value≪10^-100^). Interestingly, the highest predictor was from the combination of all four link types and DNA methylation (Fig. 6). In addition, a combined predictor using recombination hotspots and DNA methylation jointly could recapitulate up to 92% of the observed recombination rate difference at long distance and 80% at medium distance within each type of functional links (Supplementary Fig. 15).

**Figure 6.**
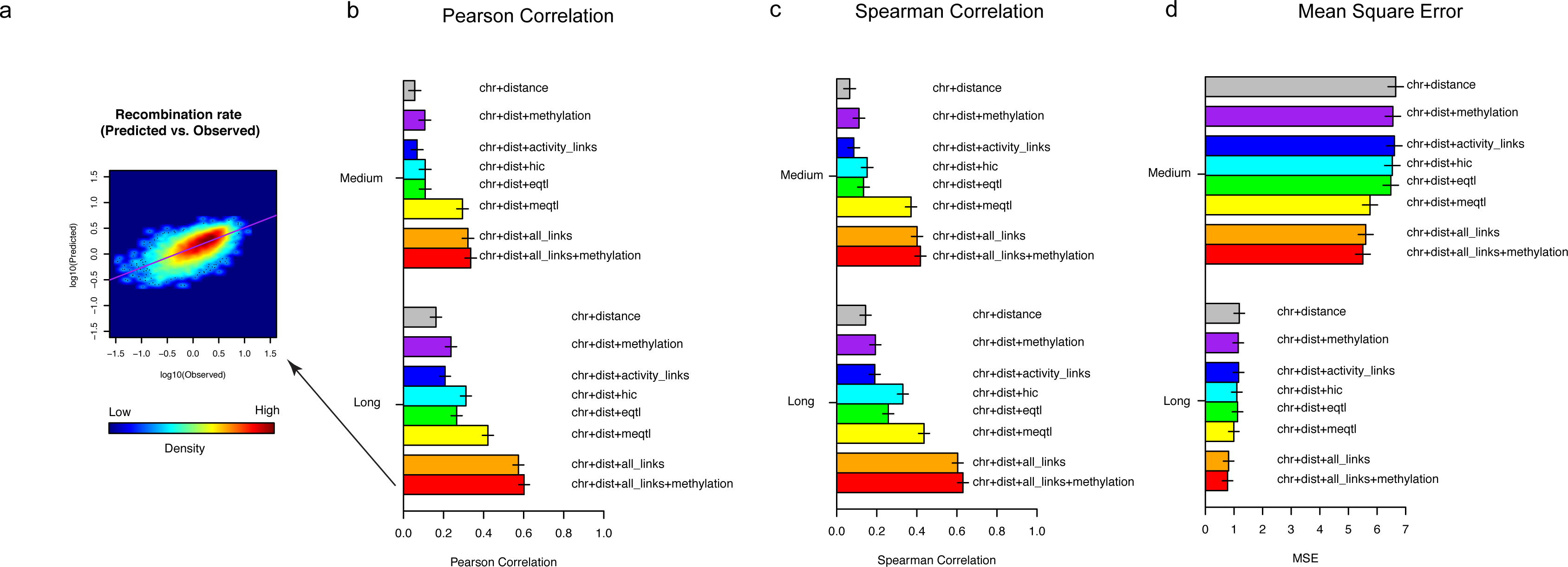
Recombination rate predictions within random intervals. (a) Heatmap of predicted recombination rate *vs*. observed recombination rate within random intervals (100kb-1Mb region, using chromosome, genomic distance, overlapped fraction with all functional links and DNA methylation level). Purple line represents the fitted linear relationship between two variables. (b) Barplot of average pearson correlation coefficient between predicted recombination rate and observed recombination rate in medium (10kb-100kb) and long distance (100kb-1Mb). (c) Barplot of average spearman correlation coefficient between predicted recombination rate and observed recombination rate in medium (10kb-100kb) and long distance (100kb-1Mb). (d) Barplot of average mean squared error (MSE) between predicted recombination rate and observed recombination rate in medium (10kb-100kb) and long distance (100kb-1Mb).

Recombination events are initiated by double-stranded breaks, which have been suggested to be associated with DNA methylation(Ha et al. 2011). Thus, DNA methylation of large regulatory domains may provide a mechanistic explanation of the reduced recombination rate. To evaluate this model, we correlated DNA methylation levels and DNA double strand break (DSB) initiation frequency both profiled within sperm. We also correlated DNA methylation levels profiled in LCL with ChIP-Seq evidence for H2A.X and gamma H2A.X, a marker of double-stranded break repair profiled in CD4+ T cell (Consortium 2012; Seo et al. 2012). Indeed, we found that increased DNA methylation showed a significant negative correlation with evidence of DNA DSB initiation frequency (Pearson −0.11, p value=1.71e^-16^) and a significant positive correlation with evidence of DNA repair (Pearson 0.36 with gamma-H2A.X, p value≪10^-100^), but a significant negative correlation with H2A.X (Pearson −0.211, p value=7.92e^-59^) (Supplementary Fig. 16). These results indicated that the increased level of DNA methylation in recombination valleys may reduce the frequency of DSB initiation and increase the efficiency of DSB repair, thus contributing to a reduced recombination rate (Fig. 5d).

## Discussion

The predominant thinking on natural selection has traditionally considered the gene as the unit of inheritance, and indeed, recombination events are depleted within gene bodies, indicating that they are inherited as intact functional units across generations. However, we expect natural selection to act simultaneously on regulatory elements and their target genes, which should be inherited as a single large unit. This would exclude potentially-disruptive recombination events between regulatory elements and their target genes which is critical for the early development stage. Consistently, we find a “recombination valley” of reduced recombination rate between regulatory elements and their target genes.

Fine-scale recombination maps across species have revealed that recombination hotspots are evolutionarily short-lived, while global patterns of recombination rate are relatively conserved(Auton et al. 2012). Our results suggest that DNA methylation could potentially drive both local and global variation in recombination rate, providing a solution to this paradox. Distantly related species have similar global DNA methylation patterns(Feng et al. 2010), suggesting the corresponding crossover rate will be relatively stable especially for the regions where the recombination hotspot density is small. However, recent genetic variation could affect DNA methylation which in turn drives local variation in recombination rate(Sigurdsson et al. 2009).

The function of global DNA demethylation in the PGC stage has remained unclear. Our results here suggest that DNA demethylation can be served as an essential event to allow recombination in the following meiosis I stage, while the high methylation level kept within functional links at oocyte stage may help the exclusion of recombination events from regulatory domains and thus keep the inherent integrity.

Together, our results indicate the evolutionary importance of transcriptional regulatory units consisting of regulatory elements and their target genes. In particular, local and global variations in recombination rate provide the important mechanistic model to interpret the relationship between genetic and epigenetic variation across individuals during the evolution. One interesting consequence is that recombination rate could be used as a prior to predict regulatory links, which could in turn be used to improve discovery of QTLs and disease associations.

## Methods

### Code availability

All scripts used in the analysis are public available at GitHub (https://github.com/dnaase/Bistools/tree/master/for_recomb2015Paper)(Lay et al. 2015). Detailed usage descriptions are elaborated in supplementary methods

## Data access

All the primary public datasets used in the analysis are shown in Supplementary Table 1.

## Acknowledgements

We thank Jianrong Wang, Pouya Kheradpour, Zhizhuo Zhang, Yongjin Park, Nezar Abdennur, Yue Li, Yu-Ping Poh, Irwin Jungreis and other members of the MIT Computational Biology group (Kellis Lab) for feedback, discussion, and suggestions. We thank Shamil R. Sunyaev for discussions and suggestions. The work was supported by the National Institutes of health via awards 1-U01-HG007610-01.

## Author contributions

Y.L. and M.K. conceived the idea. Y.L. and A.S designed and implemented the statistical test. Y.L. and M.K. performed the data analysis. Y.L., A.S. and M.K. wrote the manuscript.

